# Spike Protein-independent Attenuation of SARS-CoV-2 Omicron Variant in Laboratory Mice

**DOI:** 10.1101/2022.02.08.479543

**Authors:** Shufeng Liu, Prabhuanand Selvaraj, Kotou Sangare, Binquan Luan, Tony T. Wang

## Abstract

Despite being more transmissible, the severe acute respiratory syndrome coronavirus 2 (SARS-CoV-2) Omicron variant was found to cause milder diseases in laboratory animals, often accompanied by a lower viral load compared to previous variants of concern. This study revealed the structural basis for a robust interaction between the receptor binding domain of the Omicron spike protein and mouse ACE2. Pseudovirus bearing the Omicron spike protein efficiently utilized mouse ACE2 for entry. By comparing viral load and disease severity among laboratory mice infected by a natural Omicron variant or recombinant ancestral viruses bearing either the entire Omicron Spike or only the N501Y/Q493R mutations in its spike, we found that mutations outside the spike protein in the Omicron variant may be responsible for the observed lower viral load. Together, our results indicated that a post-entry block to the Omicron variant exists in laboratory mice.

## Main Text

The emergence of SARS-CoV-2 Omicron variant (B.1.1.529) has drastically changed the landscape of the coronavirus disease of 2019 (COVID-19) pandemic (1). Omicron appears to be displacing the Delta variant in the U.S. and possibly the rest of the world. With 30+ changes in its spike protein, the Omicron variant has been shown to evade vaccine- or natural infection-elicited immunity (2, 3). Within its receptor binding domain (RBD), the Omicron variant harbors several amino acid substitutions, including S417N, T478K, E484A, Q493R, G496S, Q498R, N501Y, Y505H, that potentially alter the species tropism of the virus. Among those, N501Y has appeared in the Alpha variant (B1.1.7), Beta (B1.351) and P.1 variant and has been demonstrated to enhance the binding of Spike to both human and mouse ACE2 (4–6). The presence of N501Y along with additional mutations (K417M, E484K, Q493R, and Q498R) is associated with mouse adaptation by serial passage (7–9). Surprisingly, several recent studies reported an attenuated phenotype in laboratory animals (10–15), raising a question whether the Omicron variant robustly infects animal species.

To tackle this question, we packaged lentiviral-based pseudoviral particles that bear no spike protein (Bald virus), the ancestral WA1 spike, and the Omicron spike protein. We then transiently transfected 293T cells with DNA plasmids to express ACE2 of 10 species. Following infection, the pseudovirus will express the green fluorescent protein (GFP). Shown in Figure 1A, the GFP+ cells were readily visible. While the majority ACE2 homologs equally mediated WA1- or Omicron-pseudovirus infection, mouse ACE2 (mACE2) expression permitted massive infection by pseudovirus bearing the Omicron spike but not the WA1 spike. Horseshoe bat ACE2, however, only mediated entry of the pseudovirus bearing the WA1 spike. Similar observations were made when using pseudoviruses carrying the firefly luciferase (FLuc) reporter gene in transient transfections or in hACE2 and mACE2 stable cell lines (Figure. 1B&C).

**Figure 1.**
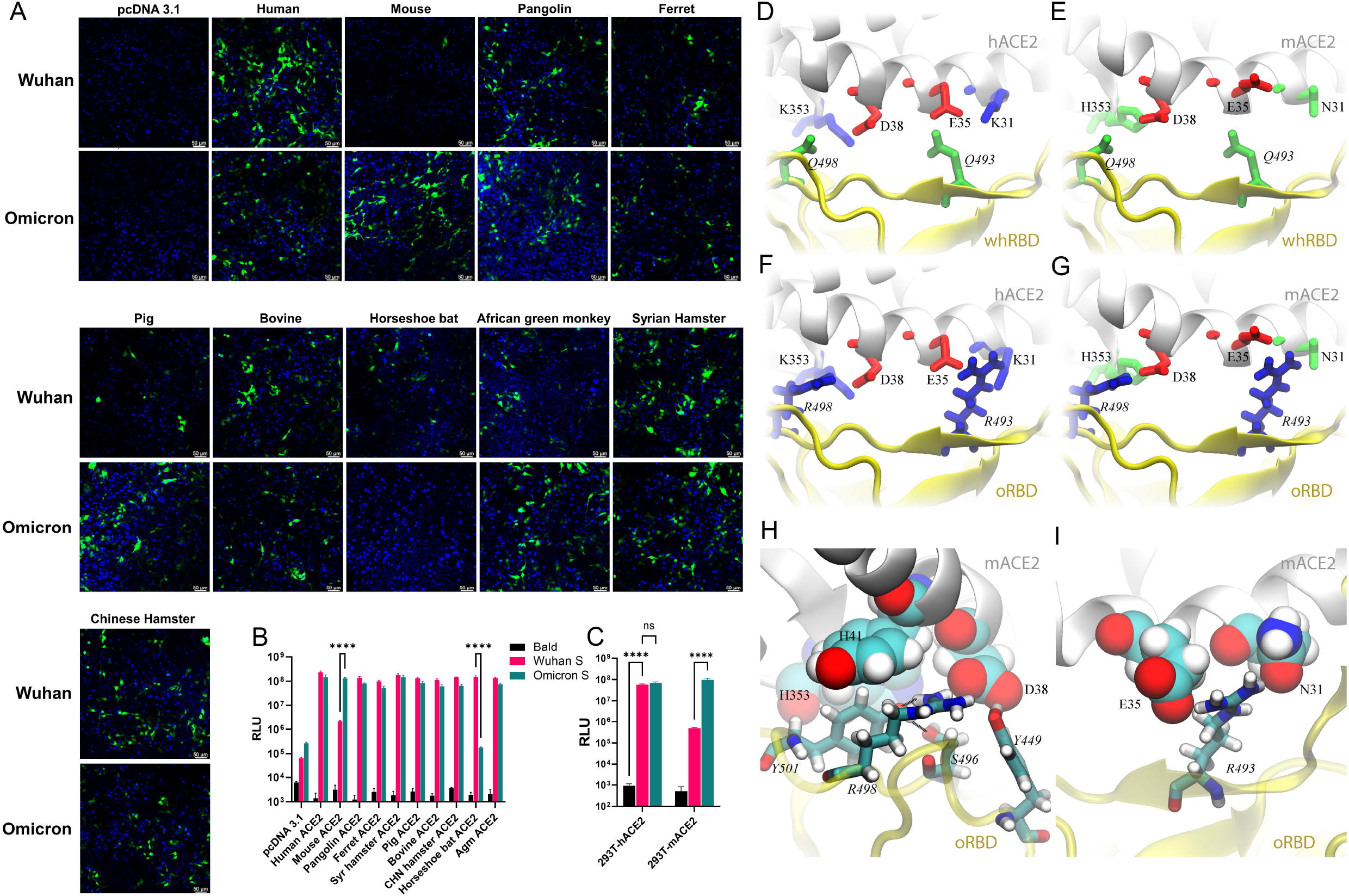
Pseudovirus bearing the Omicron spike protein efficiently infects cells expressing mACE2. (A) 293T cells were transfected with ACE2 expression plasmids of 10 species and then infected by lentiviral pseudoparticles bearing the spike protein of either the Wuhan SARS-CoV-2 variant or the Omicron variant. Infected cells would be GFP positive (green). Blue dye-stained nuclei. (B) A similar experiment was performed as in A except pseudoviruses expressing firefly luciferase were employed. (C) Infection of hACE2- and mACE2 stable 293T cells by pseudoviruses bearing the spike protein of either the Wuhan SARS-CoV-2 variant or the Omicron variant. (D—G) Various coordinations of charged residues at the ACE2-RBD interface. (D) hACE2-wtRBD (buried charge: 0), (E) mACE2-wtRBD (buried charge: −2 e), (F) hACE2-oRBD (buried charge: +2 e), (G) mACE2-oRBD (buried charge: 0). (H and I) Detailed interfacial coordinations between mACE2 and oRBD. (H) The interfacial binding near the salt-bridge R498-D38. (I)The interfacial binding near the salt-bridge R493-E35.

To understand the structural basis of mACE2-Omicron RBD (oRBD) interaction, we performed all-atom molecular dynamics (MD) simulations at physiology-like environment. Compared to the wide-type SARS-CoV-2, the Omicron variant contains 15 mutations in RBD (Figure S1A) and many of them, including the two charged mutations Q493R and Q498R, are located inside the receptor-binding motif (RBM) that directly contacts ACE2. Generally, charged residues significantly impact the protein-protein binding interface. Shown in Figure 1D, two salt-bridges (K353-D38 and E35-K31) reside at the binding surface of hACE2 while two neutral residues Q493 and Q498 are present on the Wuhan-like ancestral RBD (whRBD) surface. Besides interfacial hydrogen bonds, the two salt-bridges are partially buried after the hACE2-whRBD binding, which is energetically favorable (16). However, on the binding surface of the mACE2, negatively charged D38 and E35 are coordinated by two neutral residues H353 and N31, resulting in an overall charged surface (Figure 1E). When whRBD engages mACE2, two negatively charged residues (D38 and E35) will inevitably be buried at the binding interface, which is energetically unfavorable because the dielectric constant of a protein media is much less than that of water (Figure 1E). Different from whRBD, two positively charged residues (R493 and R498) are found in the RBM of oRBD. When hACE2 is bound with oRBD as shown in Figure 1F, the net charge at the interface is +2 e because both R493 and R498 are buried (partially) at the interface. This interfacial arrangement is energetically unfavorable and may weaken the interfacial binding, but the highly favorable N501Y mutation strengthens oRBD-hACE2 interaction. Importantly, oppositely charged oRBD and mACE2 surfaces can electrostatically attract each other, and the complex features two salt bridges (R493-E35 and R498-D38) buried at the interface with zero net charge (Figure 1G), which is energetically favorable. In support, a recent study reported mACE2 binds oRBD much better than RBDs of other variants (17).

MD simulation of the mACE2-oRBD complex in the 0.15 M NaCl electrolyte also revealed an essential atomic coordination at the interface. Figure 1H illustrates that besides forming the saltbridge with D38 in mACE2, R498 also forms a hydrogen bond with Y501, which stabilizes the conformation of Y501. Additionally, S496 forms a hydrogen bond with Y501. The latter further forms the T-shape π-πstacking with H41 (mACE2) and another parallel one with H353 of mACE2. Note that the similar interaction between Y501 and H41 was also found in other ACE2-RBD complexes, such as the hACE2 and the RBD of the Alpha variant (18), and is responsible for the enhanced interfacial interaction. Hence, in the case of mACE2-oRBD interaction, Y501 is in a locked conformation due to favorable interactions with H41 (mACE2), Y353 (mACE2), R498 (oRBD) and S496 (oRBD). Additionally, the salt-bridge between E35 (mACE2) and R493 (oRBD) further enhances the local interfacial binding (Figure 1I).

Altogether, the MD simulation implies that a buried charge inside the protein-protein interface reduces the binding affinity while a buried salt-bridge enhances the interfacial binding-affinity. To experimentally verify this hypothesis, we made several constructs expressing WA1 Spike bearing the following combinations of mutations: N501Y-Q493K, N501Y-Q493R, G496K-Q493K, N501Y-S494R, N501Y-G496K, N501F-G496K, N501K-Q493K. These mutations are expected to strengthen the RBD-mACE2 interaction through inducing salt bridges between K501/K496 of the RBD and D38 of mACE2 or between K493/R494 of RBD and E35 of mACE2. Additionally, a WA1 spike bearing Q493K/Q498Y/P499T triple mutations, which were found in a mouse-adapted virus, was included as positive control (19). Pseudoviruses bearing these spike proteins all infected 293T-mACE2 by at least two logs of magnitude compared to the WA1 spike (Figure S1B&C). These findings confirmed the flexibility of RBD-ACE2 interface in controlling species tropism. Notably, The Omicron spike protein and WA1 spike bearing Q493R/N501Y were as fusogenic as the ancestral WA1 spike in a cell-cell fusion assay (Figure S1D).

We subsequently characterized the infection of 18-month-old Balb/c mice with the Omicron variant (Isolate hCoV-19/USA/MD-HP20874/2021). Because of reported attenuation of Omicron in Syrian hamster and K18-transgenic mice (10, 11, 20), we also included a recombinant ancestral virus (WA1) bearing either the entire Omicron Spike (WA1-Omicron-S) or Q493R/N501Y (WA1-Q493R/N501Y) (Figure 2A). WA1-Omicron-S differs from the natural Omicron in that the former contains the WA1 backbone except the Omicron spike protein. Therefore, WA1-Omicron-S should enter cells in the same manner as Omicron does. If the natural Omicron variant fails to infect cells in the lung due to defective entry, as implied in several studies (21, 22), infection by the WA1-Omicron-S will similarly result in low viral load in the lung because the same entry blockage should exist. If the attenuated phenotype of the natural Omicron variant is due to poor replication or rapid elimination by host innate immune response, one might expect the WA1-Omicron-S virus would robustly infect mouse lungs. WA1-Q493R/N501Y was included in the study as a positive control because this recombinant virus efficiently infects immunocompetent mice and causes weight loss with an inoculum of 5×10^4^ pfu (Figure S2). Notably, our natural Omicron variant stock had a relatively low infectious titer (~2×10^5^ pfu/ml), we therefore chose 1×10^4^ PFU in 50 μl for intranasal inoculation in this study. Under this challenge dose, the only group of mice had modest weight loss was those inoculated with the WA1-Q493R/N501Y virus (Figure 2B). WA1-Omicron-S and Q493R/N501Y-infected mice nonetheless had clinical signs of illness at 3- and 4-day post-infection (DPI) (Figure 2C). The presence of subgenomic RNA (sgmRNA) of the E gene and total viral RNA was readily detectable in both the nasal turbinates and the lungs at 2- and 4-DPI in all three groups, although viral loads detected from WA1-Omicron-S- and WA1-Q493R/N501Y-infected mice were at least two logs higher than those from the natural variant infected animals (Figure 2D-G). The infection by WA1-Omicron-S- and WA1-Q493R/N501Y also markedly upregulated the interferon-stimulated gene 15 (ISG15) (Figure 2H) and induced pathology in the lungs (Figure 2I). Interestingly, histopathology examination of the natural Omicron variant infected lungs found that most infected animals had a low degree of immune cell infiltration (Figure 2J-K). Occasionally mice infected with the natural Omicron variant displayed more pronounced perivascular infiltrates (Figure 2L-M), despite very little viral RNA detected in the lung by RNAscope (Figure 2N-O). Mice infected by WA1-Omicron-S and WA1-Q493R/N501Y exhibited more diffuse peribronchiolar, perivascular, and alveolar inflammatory infiltrates, and with clear presence of viral RNA as detected by RNAscope using a probe that specifically targets the S gene (Figure 2P-W). The viral RNA was detected not only along bronchial epithelia but also in the lung parenchyma from WA1-Omicron-S- and WA1-Q493R/N501Y-infected mice. At 7 DPI, the histopathological changes have mostly resolved in the lungs of all three groups.

**Figure 2.**
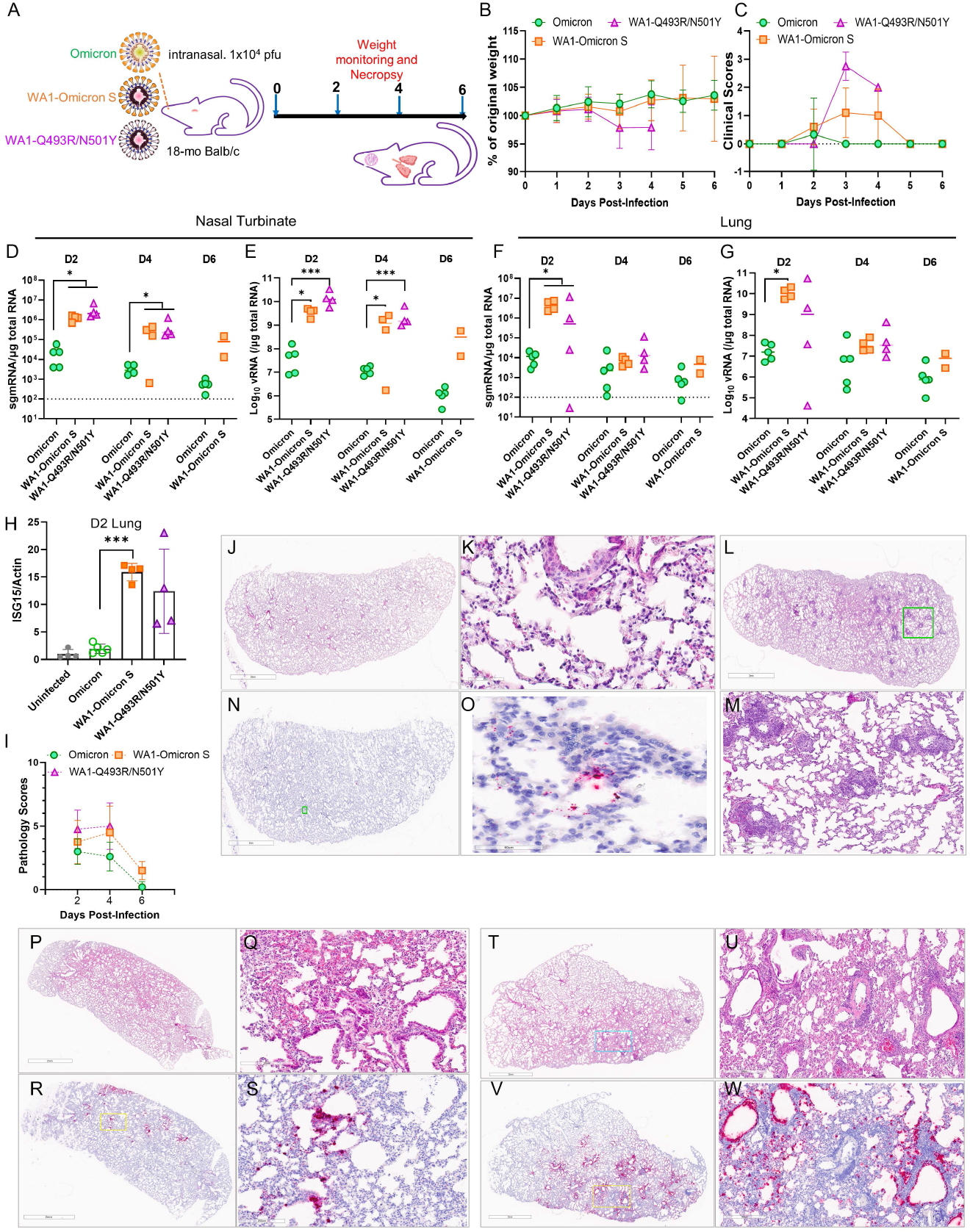
Spike-independent attenuation of the Omicron variant in Balb/c mice. (A) Overall study design. (B) Weight loss profile. (C) Clinical scores of all three groups of mice overall the period of study. Subgenomic RNA (sgmRNA) and total viral RNA levels in nasal turbinates (D-E) and lungs (F and G). (H) Induction of ISG15 at 2 DPI in mouse lungs. (I) Summed pathology scores of all three groups of mice overall the period of study. (J-M) Representative HE images of the natural Omicron infected mice. (K) is a closeup image from (J) and (M) is a closeup from (L). (N) an RNAscope image of the natural Omicron variant infected mouse, (O) is a closeup from (N). Blue, nuclei; red, viral RNA. (P) A representative HE image of the WA1-Omicron S infected mouse, (Q) is a closeup from (P). (R) a representative RNAscope image of the WA1-Omicron S infected mouse, (S) is a closeup from (R). (T) A representative HE image of the WA1-Q493R/N501Y-infected mouse, (U) is a closeup from (T). (V) a presentative RNAscope image of the WA1-Q493R/N501Y-infected mouse, (W) is a closeup from (V).

Together, these results suggest that despite an efficient usage of mACE2 by the Omicron variant, genetic changes outside the Spike protein in the natural Omicron variant may contribute to the observed attenuation in laboratory mice. Our findings argue that the low viral load detected in Omicron-infected mice is unlikely caused by an inability of the variant to enter mouse tissues, although the disease severity may still be influenced by mutations within the spike protein. The P681R mutation of the Delta variant spike protein has been linked to enhanced fusogenicity and pathogenicity in Syrian hamsters (23). Similarly, the Omicron spike is less fusogenic compared to that of the Delta variant (24, 25), but is as fusogenic as the ancestral spike protein (Figure S1D). One limitation of the current study is the relatively low inoculum and that only one natural Omicron variant was tested in this study. The low inoculum could have resulted in an overall mild disease among the three study groups. Nonetheless, findings from this study directly challenge the notion that somehow SARS-CoV-2 Omicron variant fails to reach mouse lungs. Instead, the observed lower viral load and attenuation in laboratory mice appear to be a result of post-entry blockage linked to mutations outside the spike protein in the Omicron variant.

## Acknowledgements

The following reagent was obtained through BEI Resources, NIAID, NIH: SARS-Related Coronavirus 2, Isolate hCoV-19/USA/MD-HP20874/2021 (Lineage B.1.1.529; Omicron Variant), NR-56461, contributed by Andrew S. Pekosz. We are grateful to Dr. P. Shi (UTMB) for the original 7 plasmids to perform SARS-CoV-2 reverse genetics. We thank Drs. K. Erlandson and C. Florence for assistance in obtaining critical reagents and Dr. C. Weiss for critical reading of the manuscript. The work described in this manuscript was supported by U.S. FDA intramural grant funds. The funders had no role in study design, data collection and analysis, decision to publish, or preparation of the manuscript.

## References

1. W. H. Organization, Classification of Omicron (B.1.1.529): SARS-CoV-2 Variant of Concern. https://www.who.int/news/item/26-11-2021-classification-of-omicron-(b.1.1.529)-sars-cov-2-variant-of-concern, 27th November 2021 (2021).

2. S. Cele et al., Omicron extensively but incompletely escapes Pfizer BNT162b2 neutralization. Nature 10.1038/s41586-021-04387-1 (2021).

3. C. v. S. Juliet R.C. Pulliam, Nevashan Govender, Anne von Gottberg, Cheryl Cohen, Michelle J. Groome, Jonathan Dushoff, Koleka Mlisana, Harry Moultrie, Increased risk of SARS-CoV-2 reinfection associated with emergence of the Omicron variant in South Africa. MedRxiv https://doi.org/10.1101/2021.11.11.21266068(2021).

4. H. Gu et al., Adaptation of SARS-CoV-2 in BALB/c mice for testing vaccine efficacy. Science 369, 1603–1607 (2020).

5. H. Shuai et al., Emerging SARS-CoV-2 variants expand species tropism to murines. EBioMedidne 73, 103643 (2021).

6. M. P. Xavier Montagutelli, Laurine Levillayer, Eduard Baquero Salazar, Grégory Jouvion, Laurine Conquet, Maxime Beretta, Flora Donati, Mélanie Albert, Fabiana Gambaro, Sylvie Behillil, Vincent Enouf, Dominique Rousset, Hugo Mouquet, Jean Jaubert, Felix Rey, Sylvie van der Werf, Etienne Simon-Loriere, Variants with the N501Y mutation extend SARS-CoV-2 host range to mice, with contact transmission, https://doi.org/10.1101/2021.03.18.436013 (2021).

7. M. S. Karen V. Kibler, Douglas Lake, Alexa J. Roeder, Masmudur Rahman, Brenda G. Hogue, Lok-Yin Roy Wong, Stanley Perlman, Yize Li, and Bertram L. Jacobs, Intranasal immunization with a vaccinia virus vaccine vector expressing pre-fusion stabilized SARS-CoV-2 spike fully protected mice against lethal challenge with the heavily mutated mouse-adapted SARS2-N501YMA30 strain of SARS-CoV-2. BioRxiv 10.1101/2021.12.06.471483 (2021).

8. A. Muruato et al., Mouse-adapted SARS-CoV-2 protects animals from lethal SARS-CoV challenge. PLoS Biol 19, e3001284 (2021).

9. J. Z. Lok-Yin Roy Wong, Kevin Wilhelmsen, Kun Li, Miguel E. Ortiz, Nicholas J. Schnicker, Alejandro A. Pezzulo, Peter J. Szachowicz, Klaus Klumpp, Fred Aswad, Justin Rebo, Shuh Narumiya, Makoto Murakami, David K. Meyerholz, Kristen Fortney, Paul B. McCray Jr., Stanley Perlman, Eicosanoid signaling as a therapeutic target in middle-aged mice with severe COVID-19. https://doi.org/10.1101/2021.04.20.440676(2021).

10. V. G. Katherine McMahan, Lisa H. Tostanoski, Benjamin Chung, Mazuba Siamatu, Mehul S. Suthar, Peter Halfmann, Yoshihiro Kawaoka, Cesar Piedra-Mora, Amanda J. Martinot, Swagata Kar, Hanne Andersen, Mark G. Lewis, Dan H. Barouch, Reduced Pathogenicity of the SARS-CoV-2 Omicron Variant in Hamsters. BioRxiv https://doi.org/10.1101/2022.01.02.474743 (2022).

11. P. J. Halfmann, Iida, S., Iwatsuki-Horimoto, K. et al., Adrianus C. M. Boon, Michael S. Diamond & Yoshihiro Kawaoka, SARS-CoV-2 Omicron virus causes attenuated disease in mice and hamsters. Nature https://doi.org/10.1038/s41586-022-04441-6 (2022).

12. R. J. W. Kathryn A. Ryan, Kevin R. Bewley, Christopher Burton, Oliver Carnell, Breeze E. Cavell, Amy Challis, Naomi S. Coombes, Kirsty Emery, Rachel Fell, Susan A. Fotheringham, Karen E. Gooch, Kathryn Gowan, Alastair Handley, Debbie J. Harris, Richard Humphreys, Rachel Johnson, Daniel Knott, Sian Lister, Daniel Morley, Didier Ngabo, Karen L. Osman, Jemma Paterson, Elizabeth J. Penn, Steven T. Pullan, Kevin S. Richards, Imam Shaik, Sian Summers, Stephen R. Thomas, Thomas Weldon, Nathan R. Wiblin, Richard Vipond, Bassam Hallis, Simon G. P. Yper Hall, Convalescence from prototype SARS-CoV-2 protects Syrian hamsters from disease caused by the Omicron variant. BioRxiv https://doi.org/10.1101/2021.12.24.474081(2021).

13. C. S. F. Rana Abdelnabi, Xin Zhang, Viktor Lemmens, Piet Maes, Bram Slechten, Joren Raymenants, Emmanuel André, Birgit Weynand, Kai Dallemier, Johan Neyts, The omicron (B.1.1.529) SARS-CoV-2 variant of concern does not readily infect Syrian hamsters. BioRxiv https://doi.org/10.1101/2021.12.24.474086 (2021).

14. R. Suzuki, Yamasoba, D., Kimura, I. et al., Kei Sato, Attenuated fusogenicity and pathogenicity of SARS-CoV-2 Omicron variant. Nature https://doi.org/10.1038/s41586-022-04462-1 (2022).

15. H. Shuai et al., Attenuated replication and pathogenicity of SARS-CoV-2 B.1.1.529 Omicron. Nature 10.1038/s41586-022-04442-5 (2022).

16. B. L. a. T. Huynh, Insights into SARS-CoV-2’s Mutations for Evading Human Antibodies: Sacrifice and Survival. J. Med. Chem. https://doi.org/10.1021/acs.jmedchem.1c00311(2021).

17. E. Cameroni, Bowen, J.E., Rosen, L.E. et al., Davide Corti, Broadly neutralizing antibodies overcome SARS-CoV-2 Omicron antigenic shift. Nature https://doi.org/10.1038/s41586-021-04386-2 (2021).

18. B. Luan, H. Wang, T. Huynh, Enhanced binding of the N5OlY-mutated SARS-CoV-2 spike protein to the human ACE2 receptor: insights from molecular dynamics simulations. FEBS Lett 595, 1454–1461 (2021).

19. S. R. Leist et al., A Mouse-Adapted SARS-CoV-2 Induces Acute Lung Injury and Mortality in Standard Laboratory Mice. Cell 183, 1070–1085 e1012 (2020).

20. Z.-W. Y. Shuofeng Yuan, Ronghui Liang, Kaiming Tang, Anna Jinxia Zhang, Gang Lu, Chon Phin Ong, Vincent Kwok-Man Poon, Chris Chung-Sing Chan, Bobo Wing-Yee Mok, Zhenzhi Qin, Yubin Xie, Haoran Sun, Jessica Oi-Ling Tsang, Terrence Tsz-Tai Yuen, Kenn Ka-Heng Chik, Chris Chun-Yiu Chan, Jian-Piao Cai, Cuiting Luo, Lu Lu, Cyril Chik-Yan Yip, Hin Chu, Kelvin Kai-Wang To, Honglin Chen, Dong-Yan Jin, Kwok-Yung Yuen, Jasper Fuk-Woo Chan, The SARS-CoV-2 Omicron (B.1.1.529) variant exhibits altered pathogenicity, transmissibility, and fitness in the golden Syrian hamster model. BioRxiv https://doi.org/10.1101/2022.01.12.476031 (2022).

21. B. Meng, Abdullahi, A., Ferreira, I.A.T.M. et al., Ravindra K. Gupta, Altered TMPRSS2 usage by SARS-CoV-2 Omicron impacts tropism and fusogenicity. Nature https://doi.org/10.1038/s41586-022-04474-x (2022).

22. J. C. B. Thomas P. Peacock, Jie Zhou, Nazia Thakur, Joseph Newman, Ruthiran Kugathasan, Ksenia Sukhova, Myrsini Kaforou, Dalan Bailey, Wendy S. Barclay, The SARS-CoV-2 variant, Omicron, shows rapid replication in human primary nasal epithelial cultures and efficiently uses the endosomal route of entry. BioRxiv https://doi.org/10.1101/2021.12.31.474653 (2021).

23. A. Saito et al., Enhanced fusogenicity and pathogenicity of SARS-CoV-2 Delta P681R mutation. Nature 10.1038/s41586-021-04266-9 (2021).

24. H. Zhao et al., SARS-CoV-2 Omicron variant shows less efficient replication and fusion activity when compared with delta variant in TMPRSS2-expressed cells. Emerg Microbes Infect 10.1080/22221751.2021.2023329, 1–18 (2021).

25. J. P. E. Cong Zeng, Panke Qu, Julia Faraone, Yi-Min Zheng, Claire Carlin, Joseph S. Bednash, Tongqing Zhou, Gerard Lozanski, Rama Mallampalli, Linda J. Saif, Eugene M. Oltz, Peter Mohler, Kai Xu, Richard J. Gumina, Shan-Lu Liu, Neutralization and Stability of SARS-CoV-2 Omicron Variant. BioRxiv https://doi.org/10.1101/2021.12.16.472934 (2021).

