## Supplemental Information for "Spike Protein-independent Attenuation of SARS-CoV-2 Omicron Variant in Laboratory Mice"

### Materials and Methods

#### Viruses and Cells

Vero E6 cell line (Cat # CRL-1586) cell line was purchased from American Type Culture Collection (ATCC) and cultured in eagle's minimal essential medium (MEM) supplemented with 10% fetal bovine serum (Invitrogen) and 1% penicillin/streptomycin and L-glutamine. Calu-3 cell line (Cat # HTB-55) was obtained from ATCC and maintained in EMEM+20%FBS.

The SARS-CoV-2 isolate hCoV-19/USA/MD-HP20874/2021 was obtained from BEI Resources, NIAID, NIH, and had been tittered on Vero E6 cells by plaque assay.

#### SARS-CoV-2 pseudovirus production and infection assay

Human codon-optimized cDNA encoding SARS-CoV-2 S glycoprotein of the Wuhan-Hu-1 isolate (NC_045512) or the B.1.1.529 variant (sequence available upon request) was synthesized by GenScript and cloned into eukaryotic cell expression vector pcDNA 3.1 between the BamHI and XhoI sites. Pseudovirions were produced by co-transfection of Lenti-X 293T cells with psPAX2, pTRIP-luc or pTRIP-GFP, and SARS-CoV-2 S expressing plasmid using Lipofectamine 3000. The supernatants were harvested at 48 and 72 h post-transfection and filtered through 0.45-µm membranes. Infection was done as previously described (1).

#### SARS-CoV-2 Spike-Mediated Cell–Cell Fusion Assay

To assess the Spike protein-mediated membrane fusion, we have previously established a quantitative cell-cell fusion assay in which 293T acceptor cells, containing a loxP-flanked STOP cassette that blocks transcription of the downstream luciferase reporter gene, were cocultured with Cre-expressing donor cells (1). In this assay, fusion between donor and acceptor cell membranes removes the STOP cassette and hence permits luciferase production. Various spike constructs were transfected into donor 293T cells and then the hACE2 plasmid was transfected into the acceptor 293T cells. Twenty-four hours later, recipient cells were mixed at a 1:1 ratio with 293T cells expressing Cre and CoV S (donor cells) to initiate cell–cell fusion. Luciferase activity was measured 48 h thereafter.

**Production of SARS-CoV-2 recombinant virus**

SARS-CoV-2 recombinant viruses containing 2019-nCoV/USA_WA1/2020 sequence or substitutions (Omicron S or Q493R/N501Y) in the Spike protein of WA1 was generated using a 7-plasmid reverse genetic system which was based on the virus strain (2019-nCoV/USA_WA1/2020) isolated from the first reported SARS-CoV-2 case in the U.S. (2). The initial 7 plasmids were generous gifts from Dr. P-Y Shi (UTMB). Upon receipt, fragment 4 was subsequently subcloned into a low-copy plasmid pSMART LCAmp (Lucigen) to increase stability. Standard molecular biology technique was employed to create mutations. In vitro transcription and electroporation were carried following procedures that were detailed elsewhere (3, 4). Recombinant viruses were further deep sequenced to confirm the presence of mutations.

**Mouse Infection Experiments**

Aged male and female Balb/c mice were previously purchased from the Jackson laboratory and held at FDA vivarium. All experiments were performed within the biosafety level 3 (BSL-3) suite on the White Oak campus of the U.S. Food and Drug Administration. The study protocol details were approved by the White Oak Consolidated Animal Care and Use Committee and carried out in accordance with the PHS Policy on Humane Care & Use of Laboratory Animals.

For infection studies, mice were first anesthetized by 3-5% isoflurane. Intranasal inoculation was done by pipetting $10^4$ PFU SARS-CoV-2 in 50 µl volume dropwise into the nostrils of the mouse. Mice were weighed and observed daily. For tissue collections, mice were euthanized by $CO_2$ overdose on days 2, 4, 6 as necessary. Categories included in clinical scoring include weight loss (0–1), posture and appearance of fur (0–1), appearance of lethargy (0–2), and eye closure (0–1).

**RNA isolation from tissues**

RNA was extracted from 0.1-gram tissue homogenates using Trizol and the RNeasy Mini kit (Qiagen) and eluted with 50 µL of water. One microgram RNA was used for each reaction in real-time RT-PCR.

**Real-time PCR assay of SARS-CoV-2 viral and subgenomic RNA**

Quantification of SARS-CoV-2 viral RNA (vRNA) was conducted using the SARS-CoV-2 (2019-nCoV) CDC qPCR Probe Assay (IDTDNA) using iTaq Universal Probes One-Step Kit (Bio-Rad). The standard curve was generated using 2019-nCoV_N_Positive Control (IDTDNA). The detection limit of the vRNA was determined to be 100 copies/reaction. Quantification of SARS-CoV-2 E gene subgenomic mRNA (sgmRNA) was conducted using Luna Universal Probe One-Step RT-qPCR Kit (New England Biolabs) on a Step One Plus Real-Time PCR system (Applied Biosystems). The primer and probe sequences were: SARS2EF: CGATCTCTTGTAGATCTGTTCT; PROBE: FAM-ACACTAGCCATCCTTACTGCGCTTCG-BHQ-1; SARS2ER: ATATTGCAGCAGTACGCACACA. To generate a standard curve, the cDNA of SARS-CoV-2 E gene sgmRNA was cloned into a pCR2.1-TOPO plasmid. The copy number of sgmRNA was calculated by comparing to a standard curve obtained with serial dilutions of the standard plasmid. The limit of quantification (LoQ) of the sgmRNA was determined to be 100 copies/reaction. Values below LoQ were mathematically extrapolated based on the standard curves for graphing purpose. When graphing the results in Prism 9, values below LoQ were arbitrarily set to half of the LoQ values.

**Histopathology Analyses**

Tissues (lungs and nasal turbinates) were fixed in 10% neural buffered formalin for 2-7 days and then processed for paraffin embedding. The 5-μm sections were stained with hematoxylin and eosin for histopathological examinations. Images were scanned using an Aperio ImageScope and scored under the following categories: consolidation, alveolar wall thickening, alveolar airway infiltrates, perivascular Infiltrates, perivascular edema, peribronchiolar infiltrates, type II pneumocyte hyperplasia, necrosis (alveoli and bronchiole), bronchiole mucosal hyperplasia, bronchiole airway infiltrates, proteinaceous fluid, hemorrhage, vasculitis. Grading scale: 0 = none, 1 = mild, 2 = moderate, 3 = severe.

**In-situ Hybridization**

To detect SARS-CoV-2 genomic RNA in FFPE tissues, ISH was performed using the RNAscope 2.5 HD RED kit, a single plex assay method (Advanced Cell Diagnostics; Catalog 322373) according to the manufacturer's instructions. Briefly, Mm PPIB probe detecting peptidylprolyl isomerase B gene (housekeeping gene) (catalog 313911, positive-control RNA probe), dapB probe detecting dihydrodipicolinate reductase gene from Bacillus subtilis strain SMY (a soil bacterium) (catalog 310043, negative-control RNA probe) and V-SARS-Cov-2-S (catalog 854841) targeting SARS-CoV-2 positive-sense (genomic) RNA. Tissue sections were deparaffinized with xylene, underwent a series of ethanol washes and peroxidase blocking, and were then heated in kit-provided antigen retrieval buffer and digested by kit-provided proteinase. Sections were exposed to ISH target probes and incubated at 40°C in a hybridization oven for 2 hours. After rinsing, ISH signal was amplified using kit-provided pre-amplifier and amplifier conjugated to alkaline phosphatase and incubated with a fast-red substrate solution for 10 minutes at room temperature. Sections were then stained with 50% hematoxylin solution followed by 0.02% ammonium water treatment, dried in a 60°C dry oven, mounted, and stored at 4°C until image analysis.

**Statistical analysis**

All experiments were independently performed at least twice as indicated in the Figure legends. Except were specified, bar graphs were plotted to show mean ± standard deviation (SD). Statistical analyses were performed using Prism 9.

**MD simulations**

We carried out all-atom MD simulations for a complex of the mouse's ACE2 and the RBD of Omicron's spike protein using the NAMD2.13 package (5) running on the IBM Power Cluster. We obtained the secondary structure of the mouse's ACE2 (mACE2) through the homology modelling according to the structure of human ACE2 (PDB code: 6M0J) using SWISS-MODEL (6). The atomic structure of the Omicron RBD (oRBD) was adopted from the recently resolved crystal structure (PDB code: 7T9L) (7). The complex of mACE2 and oRBD was assembled by the structural alignment to the known complex structure of hACE2 and oRBD (PDB code: 7T9L). After solvating the mACE2-oRBD complex in a 0.15 NaCl electrolyte, we performed all-atom molecular dynamics simulations to equilibrate the built complex in its physiology-like environment. The final simulation system comprises 209,460 atoms.

We used the CHARMM36m force field (8) for proteins, the TIP3P model(9, 10) for water, the standard force field (11) for $Na^+$ and $Cl^-$. The periodic boundary conditions (PBC) were applied in all three dimensions. Long-range Coulomb interactions were computed using particle-mesh Ewald (PME) full electrostatics with the grid size of about 1 Å in each dimension. The pair-wise van der Waals (vdW) energies were calculated using a smooth (10-12 Å) cutoff. The temperature *T* was kept at 300 K by applying the Langevin thermostat (12),while the pressure was maintained constant at 1 bar using the Nosé-Hoover method (13). With the SETTLE algorithm (14) enabled to keep all bonds rigid, the simulation time-step was 2 fs for bonded and non-bonded (including vdW, angle, improper and dihedral) interactions, and the time-step for Coulomb interactions was 4 fs, with the multiple time-step algorithm (15).

### References


1. S. Liu *et al.*, The PRRA insert at the S1/S2 site modulates cellular tropism of SARS-CoV-2 and ACE2 usage by the closely related Bat RaTG13. *J Virol* 10.1128/JVI.01751-20 (2021).
2. X. Xie *et al.*, An Infectious cDNA Clone of SARS-CoV-2. *Cell Host Microbe* **27**, 841-848 e843 (2020).
3. X. Xie *et al.*, Engineering SARS-CoV-2 using a reverse genetic system. *Nat Protoc* **16**, 1761-1784 (2021).
4. T. H. Shufeng Liu, Charles B Stauft, Tony T Wang, Binquan Luan, Structure-Function Analysis of Resistance to Bamlanivimab by SARS-CoV-2 Variants Kappa, Delta, and Lambda *J Chem Inf Model* **61**, 5133-5140 (2021).
5. J. C. Phillips *et al.*, Scalable molecular dynamics with NAMD. *J Comput Chem* **26**, 1781-1802 (2005).
6. A. Waterhouse *et al.*, SWISS-MODEL: homology modelling of protein structures and complexes. *Nucleic Acids Res* **46**, W296-W303 (2018).
7. D. Mannar *et al.*, SARS-CoV-2 Omicron variant: Antibody evasion and cryo-EM structure of spike protein-ACE2 complex. *Science* 10.1126/science.abn7760, eabn7760 (2022).
8. J. Huang *et al.*, CHARMM36m: an improved force field for folded and intrinsically disordered proteins. *Nat Methods* **14**, 71-73 (2017).
9. W. L. C. Jorgensen, J.; Madura, J. D.; Impey, R. W.; Klein, M. L. , Comparison of Simple Potential Functions for Simulating Liquid Water. *J. Chem. Phys.* **79**, 926–935 (1983).
10. E. F. Neria, S.; Karplus, M., Simulation of Activation Free Energies in Molecular Systems. *J. Chem. Phys.* **105**, 1902–1921 (1996).
11. D. R. Beglov, B, Finite representation of an infinite bulk system: Solvent boundary potential for computer simulations. *J. Chem. Phys.* **100**, 9050–9063 (1994).
12. M. P. T. Allen, D. J, Computer Simulation of Liquids. *Oxford University Press: New York* (1987).
13. G. J. T. Martyna, D. J.; Klein, M. L, Constant Pressure Molecular Dynamics Algorithms. *J. Chem. Phys.* **101**, 4177–4189 (1994).
14. S. K. Miyamoto, P. A. , SETTLE: An Analytical Version of the SHAKE and RATTLE Algorithm for Rigid Water Molecules. *J. Comp. Chem* **13**, 952–962 (1992).
15. M. B. Tuckerman, B. J.; Martyna, G. J. , Reversible multiple time scale molecular dynamics. . *The Journal of chemical physics* **97**, 1990–2001 (1992).

**Supplemental Figure Legends**

**Figure S1. The spike protein of Omicron variant carries mutations that confer utility of mouse ACE2 for entry**. (A) Illustration of all changes found in the Omicron isolate used in this study (hCoV-19/USA/MD-HP20874/2021) in comparison to the USA-WA1/2020. Fortuitously, the Omicron spike protein in pseudoviruses also carries the identical number of changes. (B) and (C) Pseudoviruses carrying WA1 spike with indicated combinations of mutations infected 293T-hACE2 (B) or 293-mACE2 cell lines. (D) Cell-cell fusion is mediated by WA1 spike or Omicron spike or the WA1-Q493R/N501Y spike protein.

**Figure S2.** A recombinant SARS-CoV-2 (i.e., WA1-Q493R/N501Y) infects laboratory mice. (A) Overall study design (B) Weight loss profile. (C) sgmRNA levels in nasal turbinates (NB) and the lungs at 2 DPI. Representative HE and RNAscope images of an entire lobe of the lung from the uninfected (D), WA1 infected (E), and WA1-Q493R/N501Y-infected mice (F). Closeup images are also included in (F).

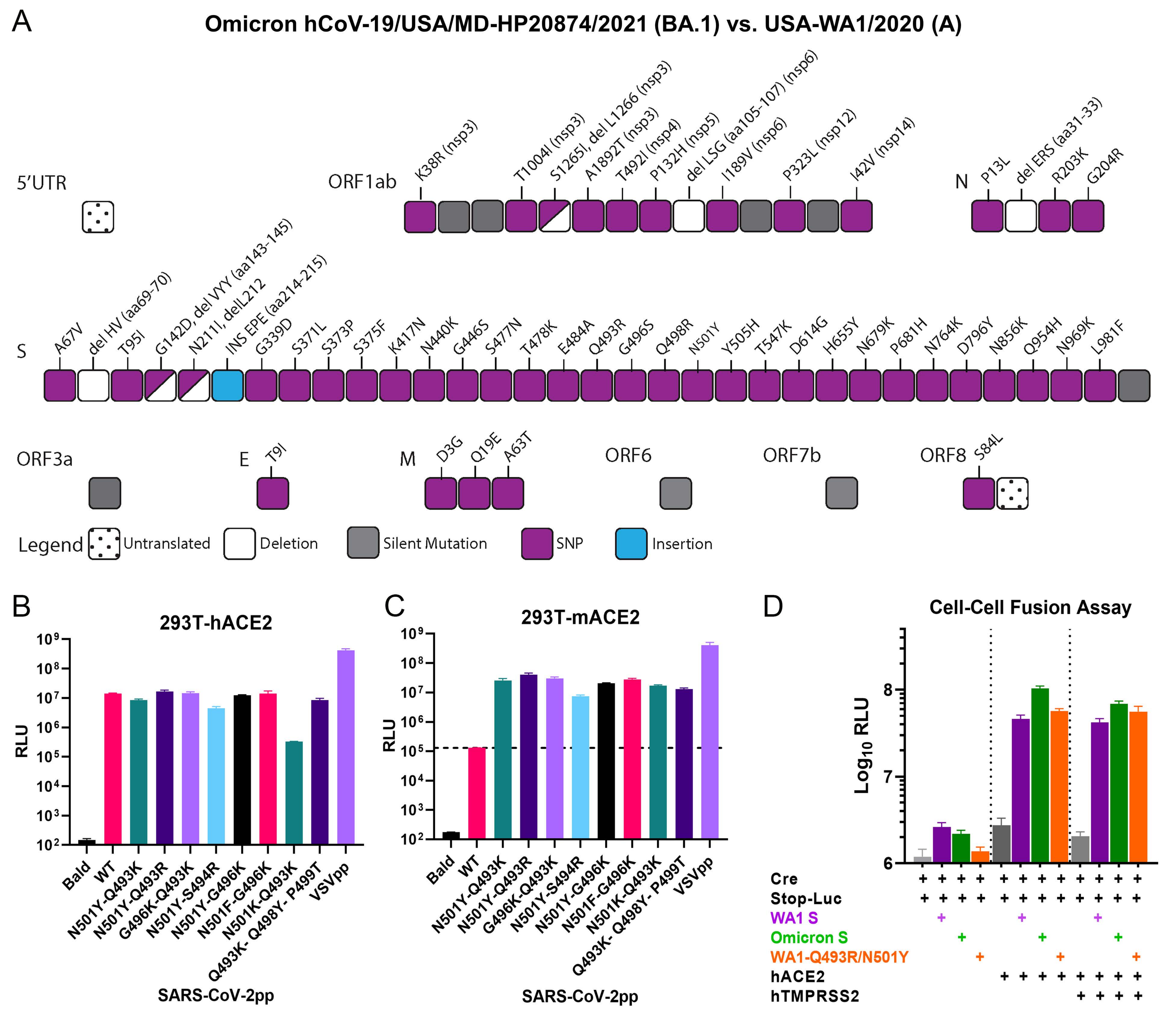
A
Omicron hCoV-19/USA/MD-HP20874/2021 (BA.1) vs. USA-WA1/2020 (A)
5'UTR
ORF1ab
K38R (nsp3)
T1004I (nsp3)
S1265I, del L1266 (nsp3)
A1892T (nsp3)
T492I (nsp4)
P132H (nsp5)
del LSG (aa105-107) (nsp6)
I189V (nsp6)
P323L (nsp12)
I42V (nsp14)
N
P13L
del ERS (aa31-33)
R203K
G204R
S
A67V
del HV (aa69-70)
T95I
G142D, del VYY (aa143-145)
N211I, delL212
INS EPE (aa214-215)
G339D
S371L
S373P
S375F
K417N
N440K
G446S
S477N
T478K
E484A
Q493R
G496S
Q498R
N501Y
Y505H
T547K
D614G
H655Y
N679K
P681H
N764K
D796Y
N856K
Q954H
N969K
L981F
ORF3a
E
T9I
M
D3G
Q19E
A63T
ORF6
ORF7b
ORF8
S84L
Legend
Untranslated
Deletion
Silent Mutation
SNP
Insertion
B
293T-hACE2
RLU
Bald
WT
N501Y-Q493K
N501Y-Q493R
G496K-Q493K
N501Y-S494R
N501Y-G496K
N501F-G496K
N501K-Q493K
Q493K- Q498Y- P499T
VSVpp
SARS-CoV-2pp
C
293T-mACE2
RLU
SARS-CoV-2pp
D
Cell-Cell Fusion Assay
Log10 RLU
Cre
Stop-Luc
WA1 S
Omicron S
WA1-Q493R/N501Y
hACE2
hTMPRSS2

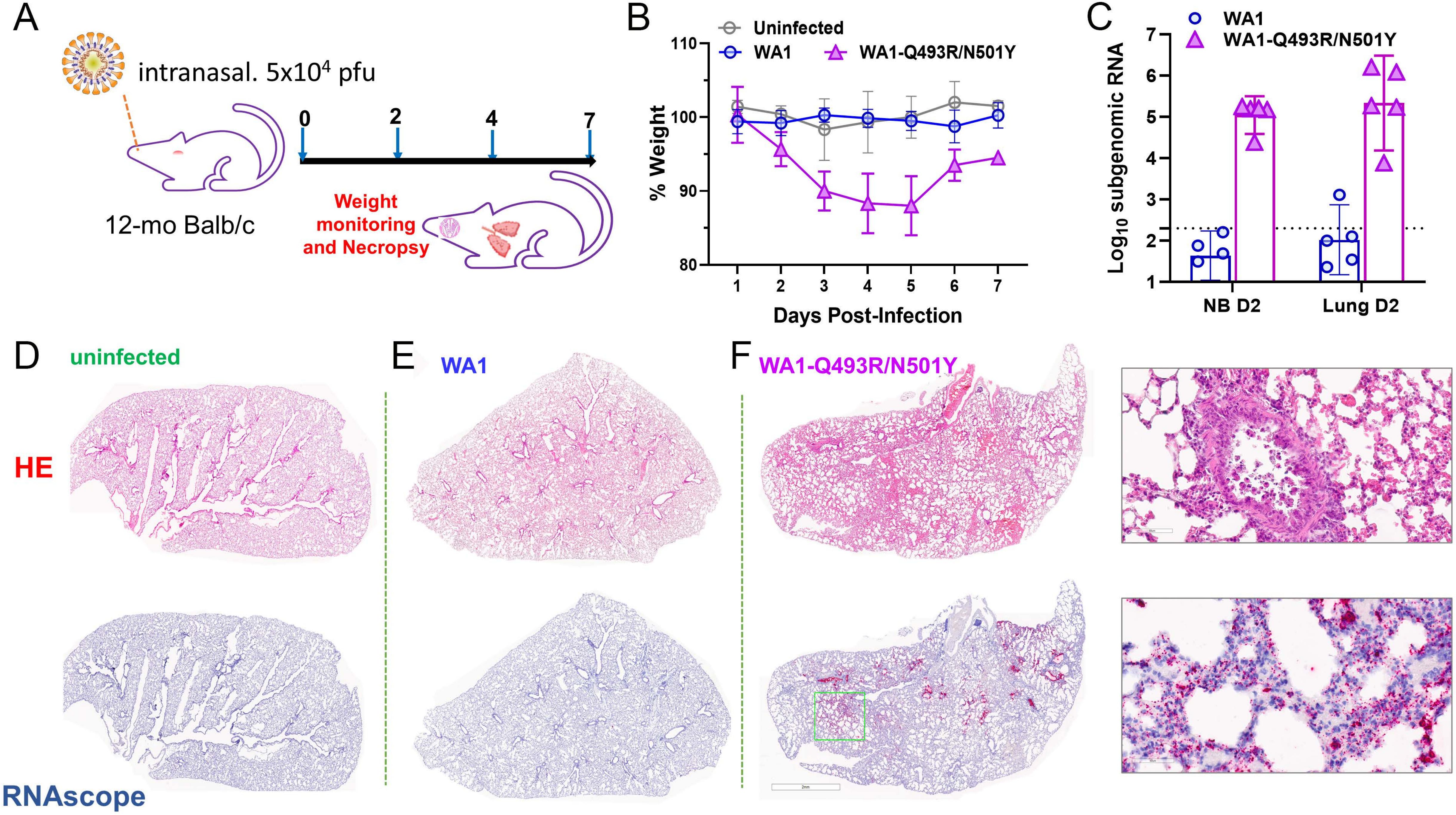

A
intranasal. 5x10^4 pfu
12-mo Balb/c
0
2
4
7
Weight monitoring and Necropsy
B
Uninfected
WA1
WA1-Q493R/N501Y
% Weight
Days Post-Infection
C
WA1
WA1-Q493R/N501Y
Log10 subgenomic RNA
NB D2
Lung D2
D
uninfected
E
WA1
F
WA1-Q493R/N501Y
HE
RNAscope
